# Tyrosine hydroxylase–mediated neuroimmune crosstalk regulates antitumor immunity in glioblastoma during oncolytic herpes virotherapy

**DOI:** 10.64898/2026.06.20.733517

**Authors:** Konstantina Kyritsi, Haocheng Ding, Dong Zhu, Ravindra Kollhe, Theodore S. Johnson, Balveen Kaur, David H. Munn, Bangxing Hong

## Abstract

Neuroimmune crosstalk is increasingly recognized as a key regulator of tumor progression and therapeutic response, yet its role in central nervous system (CNS) tumors remains poorly understood. Here, we investigate tyrosine hydroxylase (TH)–mediated neuronal signaling in glioblastoma (GBM) and its impact on antitumor immunity and response to oncolytic virotherapy (OV). We show that TH⁺ cells are widely distributed within the GBM microenvironment, including neurons, astrocytes, and immune cells, and are enriched at the tumor margin. In addition, TH⁺ cells are present in the tumor-draining lymph nodes (TDLNs) of GBM, where they localize near lymphatic vessels and are associated with lymphangiogenesis. Notably, a subset of CD3⁺TH⁺ T cells is detected within lymphatic structures of TDLNs, suggesting immune-intrinsic catecholamine signaling. Single-cell RNA sequencing reveals that noradrenergic signaling, particularly via β2-adrenergic receptors (ADRB2), predominates in tumor-infiltrating myeloid cells and is dynamically regulated by therapy. Intratumoral administration of oncolytic herpes simplex virus (oHSV) upregulates ADRB2 expression in macrophages, whereas systemic chemo-immunotherapy induces distinct receptor modulation patterns in tumors and TDLNs. Functionally, pharmacologic β-adrenergic blockade significantly enhances the efficacy of oHSV therapy in orthotopic GBM and subcutaneous melanoma models, resulting in reduced tumor growth, increased tumor cell death, and enhanced CD8⁺ T cell infiltration. Similarly, direct inhibition of TH enzymatic activity suppresses tumor progression and further potentiates OV. Mechanistically, TH inhibition not only promotes tumor-infiltrated cytotoxic immune cells CD8, NK and γδ T cells, but also suppresses the activity of immunosuppressive myeloid cells, including transcriptional (Fos), and metabolism (Arg) modification in M2 macrophages and other immune cells. Collectively, these findings identify TH-mediated neuroimmune signaling as a critical regulator of tumor immunity in GBM and demonstrate that targeting catecholaminergic pathways or downstream neuroimmune crosstalk pathways can enhance the efficacy of OV. This study provides a rationale for integrating neural modulation into immunotherapeutic strategies for CNS malignancies.

## Introduction

Neuroimmune crosstalk has emerged as a critical regulator of tumor progression and therapeutic response in solid malignancies^1–7^. In peripheral tissues, interactions between the peripheral nervous system (PNS) and immune cells have been increasingly recognized as key modulators of tumor growth^8,9^, metastasis^10^, treatment failure^10–14^ and antitumor immunity^11^. However, the role of neuron–immune interactions in central nervous system (CNS) tumors remains poorly understood. This gap in knowledge is particularly significant given the unique anatomical and immunological features of the CNS tumor microenvironment.

Glioblastoma (GBM), the most aggressive primary brain tumor, develops within a highly innervated neural environment composed of diverse neuronal subtypes^2,3,5–7^. Among these, tyrosine hydroxylase (TH)-expressing neurons^15^—key regulators of catecholamine synthesis—represent a major neuronal population capable of directly interacting with tumor cells and immune components in GBM. Emerging evidence suggests that TH expression is not restricted to neurons but may also be detected in non-neuronal immune cells, including CD4 T-cells^16^ and macrophages^17,18^, further complicating its role in neuroimmune regulation. Through modulation of neurotransmitter signaling, TH-expressing cells may influence immune cell recruitment, activation, and function, thereby impacting antitumor immunity.

Oncolytic virotherapy has shown promise as a novel therapeutic strategy for GBM by selectively infecting and lysing tumor cells while simultaneously stimulating antitumor immune responses^19,20^. However, the extent to which neuroimmune signaling—particularly TH-mediated pathways—modulates the efficacy of virotherapy remains unclear. In addition, how local virotherapy influences neuronal signaling within both the tumor and tumor-draining lymph nodes (TDLNs) has not been systematically investigated.

In this study, we characterize the role of TH⁺ signaling in GBM and investigate how oncolytic herpes virotherapy (oHSV) reshapes TH-mediated neuroimmune crosstalk in the tumor microenvironment. Our findings provide new insights into the interplay between the nervous and immune systems in CNS tumors and identify potential therapeutic opportunities to enhance the efficacy of immunovirotherapy.

## Materials and Methods

### Cell lines, oncolytic virus and reagents

Mouse GSC005 were cultured as spheres in Dulbecco’s Modified Eagle Medium (DMEM)/F12 media (Sigma-Aldrich D80620) supplemented with 2 mM L-glutamine, 1% N2 supplement, 0.5% penicillin streptomycin, recombinant human epidermal growth factor (EGF) (20 ng/mL), recombinant human FGF-basic (20 ng/mL) and puromycin. Accutase was used for spheres dissociation and passaging. B16 cells are cultured in DMEM added 10 % fetal bovine serum (FBS), and 1% Penicillin/Streptomycin. All cells were incubated at 37^0^C, validated to lack of contamination and maintained below thirty passages before the experiment started. All cells tested and confirmed negative for mycoplasma (Mycoplasma PCR Detection Kit, #G238, Applied Biological Materials, BC, Canada). β-blocker (propranolol) and TH inhibitor (α-Methyl-p-tyrosine, AMPT,THi) were purchased from MedChemExpress. oHSV used igenes anddy is PKR-shRNA virus (oHSV-shPKR and oHSV-mshPKR) generated in the lab^21^, which contains double mutation of ICP6 and gamma 34.5 genes, and expresses PKR-shRNA to increase virus infection and replication capacity in tumor cells.

### Animal models

Animal experiments were performed consonantly with protocols approved by Augusta University (protocol #: 2023-1098). C57/BL6 (JAX:000664) mice were purchased from the Jackson Laboratory, Bar Harbor, ME. All mice, five per cage, were housed in room-controlled temperature on a 12:12-h light cycle with access to food, standard chow pellet diet, and water. Before the mice underwent stereotactic intracranial injection, anesthesia was performed with intraperitoneal injection of 0.2 mL of stock solution containing the following drugs: ketamine HCl (25 mg/mL) and xylazine (2.5 mg/mL) diluted in distilled water. Mice were fixed in a stereotactic device (David Kopf Instruments, Tujunga, CA), the surgical site anointed with 70% ethyl alcohol and a skin incision was performed over the midline. Over the right hemisphere at a location 2 mm lateral and 1 mm anterior to bregma, GSC005 cells or oHSV in 2 μL PBS were intra-cranially injected using needles (Hamilton 80300 for cell implantation and Hamilton 80000 for virus) at a depth of 3.5 mm and at a rate of 0.4 μL/min using autoinjectors (KD Scientific Inc, Holliston, MA). Slowly, needles were removed, and skin was sutured using a 5-0 nylon thread. Seven days after tumor cells injection, mice were treated intra-tumorally with sterile PBS or 2 x 10^5^ PFU of oHSV and started daily treatment by intraperitoneal injection with β-blocker (2 mg/kg/day) or TH inhibitor (25 mg/kg/day) until analysis. Subcutaneous B16 model was established in C57BL/6J mice. Animals were observed and euthanized at the indicated time points and brains were harvested.

### Immunofluorescence and immunohistrochemistry staining

Mouse 005 and B16 tumors were collected to prepare paraffin blocks and sections. After deparaffinization and antigen retrieval, the sections were permeabilized with 0.04% triton-X and blocked with 2% goat serum and incubated with primary antibodies (TH, Lyve1, NeuN, GFAP, CD8, cleaved caspase-3, Ki-67) overnight at 4^0^C. The day after, the sections were washed and incubated with secondary antibodies for 1 hour. Slides were fixed with cover slips using Fluor mount-G^TM^, with DAPI (Invitrogen, E142914) solution, images were taken by fluorescence microscopy (Nikon Eclipse Ts2) and analyzed using ImageJ software.

### Single-Cell mRNA sequencing

CD45+ cells were isolated from tumors treated with, oHSV, or oHSV and TH inhibitor. mRNA libraries were constructed using the 10x Genomics Chromium Next GEM Single Cell 5’ HT Reagent Kits v2 (Dual Index). Raw sequencing data files were demultiplexed into FASTQ files and analyzed using the Cell Ranger algorithm (10x Genomics). The filtered count matrices and filtered contig V(D)J annotations were analyzed with R (v 4.2) using Seurat and Bioconductor packages. The low-quality cells were filtered out retaining cells with detected gene numbers >200, and mitochondrial genes <15%. Genes that were expressed by less than 3 cells were rejected.

### Bioinformatics Analysis

Analysis of dopaminergic receptors and adrenergic receptors expression from human GBM tumors and brain cells were assed using data from GBMseq (www.gbmseq.org)^22^.

### Statistical analysis

All quantitative results are presented as means ± standard deviation (SD). Statistical differences between two groups were assessed using the Mann-Whitney U test or Student’s *t*-test. For comparisons involving more than two groups, ANOVA was employed. Statistical analyses were conducted using Prism 5 software (GraphPad Software, Inc., La Jolla, CA). A p-value of less than 0.05 was considered statistically significant.

## Results

### TH-expressing neuronal and non-neuronal cells are present within GBM tumors and at the tumor margins

TH is the rate-limiting enzyme in the biosynthesis of catecholamine neurotransmitters, including dopamine, norepinephrine, and epinephrine^23^. In the PNS, TH⁺ neurons are predominantly sympathetic neurons. In contrast, within the CNS, TH⁺ neurons primarily consist of dopaminergic and noradrenergic neuronal populations, whereas adrenergic neurons are relatively rare^24^. Notably, TH expression is not restricted to neuronal cells; it has also been detected in non-neuronal populations, including immune cells, where it can regulate cellular function^16^.

Immunohistochemical (IHC) staining revealed that TH⁺ neuronal cell bodies are detectable in adjacent normal brain tissue in murine glioblastoma GSC005 tumor–bearing mice (**Fig. 1a**) and are highly enriched at the brain–tumor interface (**Fig. 1b**). To further characterize TH⁺ cells within GBM in syngeneic mouse models, co-immunostaining was performed using TH together with the neuronal marker NeuN and the astrocyte marker GFAP. These analyses demonstrated the presence of TH⁺ neurons (**Fig. 1c**) as well as TH⁺ astrocytes (**Fig. 1d**) within the tumor microenvironment.

**Fig. 1.**
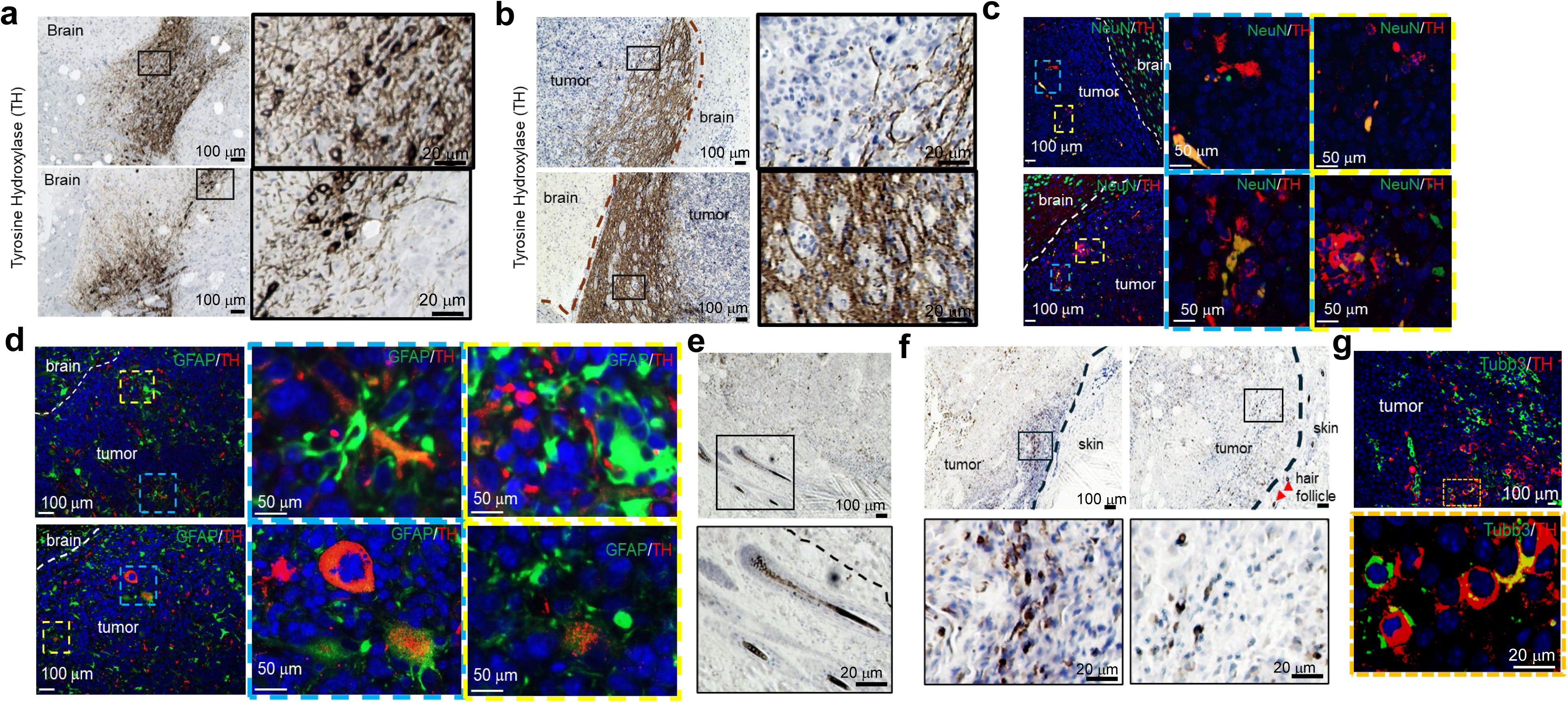
TH expression in GBM and melanoma. (**a-b**). Immunohistochemistry staining of TH in adjacent brain parecnhymal (**a**) and GSC005 tumor (**b**) from C57BL/6J tumor-bearing mice. Photos shown one of the representative staining. (**c-d**). Immunofluorescence staining of TH and NeuN (**c**) and TH and GFAP (**d**) of GSC005 tumor from C57BL/6J mice. Photos shown one of the representative staining. (**e-f**). Immunohistochemistry staining of TH in adjacent normal skin (**e**) and B16F10 tumor (**f**) from C57BL/6J tumor-bearing mice. Photos shown one of the representative staining. (**g**). Immunofluorescence staining of TH and Tubb3 of GSC005 tumor from C57BL/6J mice. Photos shown one of the representative staining.

TH⁺ sympathetic neurons in the PNS have been reported to regulate the exhaustion of tumor-infiltrating T cells^11^. To compare TH-mediated signaling between CNS tumors and non-CNS solid tumors innervated by the PNS, a subcutaneous B16 melanoma model was employed in the study. In this model, TH expression was detected not only in tumor-adjacent skin structures, such as hair follicles (**Fig. 1e**), but also within the tumor tissue itself (**Fig. 1f**). Further analysis indicated that these TH⁺ cells comprise both neuronal and non-neuronal populations (**Fig. 1g**).

### TH signaling regulates tumor-draining lymph nodes

TH⁺ neuronal signaling not only directly influences tumor development but also modulates immune cell function within the tumor microenvironment^11^. In addition, TH⁺ neurons have been reported to innervate lymph nodes^25^, suggesting a broader role in regulating systemic immune responses. In the context of central nervous system (CNS) tumors, the mechanisms of antigen presentation remain incompletely understood. However, the deep cervical lymph nodes (dcLNs) are widely recognized as the primary tumor-draining lymph nodes (TDLNs) that mediate antitumor immune responses in GBM^26^.

To investigate potential differences in TH⁺ innervation and lymphatic drainage between ipsilateral and contralateral dcLNs in GSC005 tumor-bearing mice, immunofluorescence staining was performed for TH and Lyve-1, a canonical lymphatic endothelial marker. The results demonstrated that TH⁺ cells are aligned in close proximity to lymphatic vessels in both ipsilateral and contralateral dcLNs (**Fig. 2a**). Notably, a marked expansion of lymphatic vessels (lymphangiogenesis) was observed in ipsilateral dcLNs compared to contralateral controls (**Fig. 2a**).

**Fig. 2.**
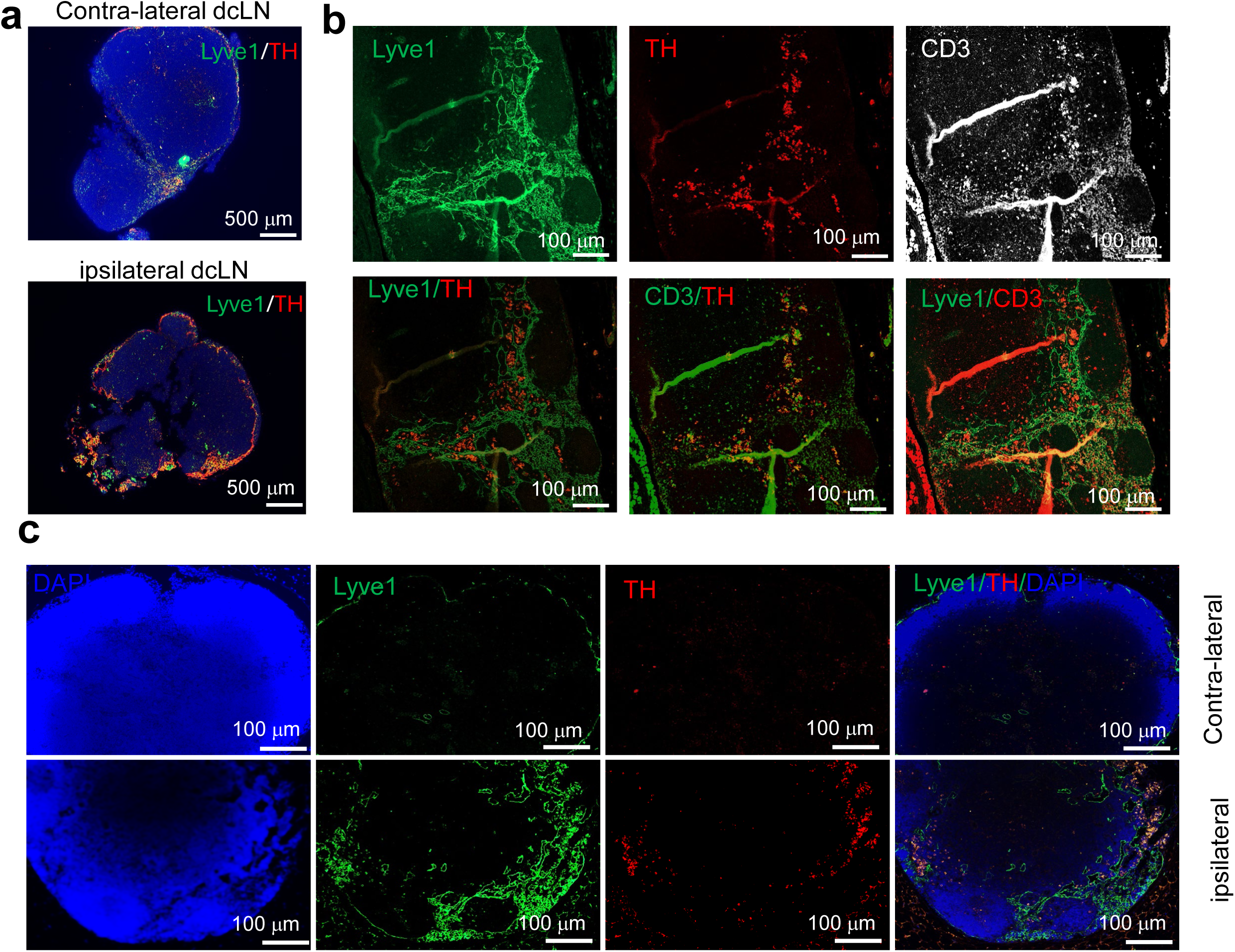
TH expression in tumor-draining lymph nodes (TDLNs). (**a**). Immunofluorescence staining of TH and Lyve1 from contra-lateral and ipsilateral deep cervical lymph nodes (dcLNs) from GSC005 tumor-bearing C57BL/6J mice. Photos shown one of the representative staining. (**b**). Immunofluorescence staining of TH, CD3 and Lyve1 from ipsilateral dcLNs from GSC005 tumor-bearing C57BL/6J mice. Photos shown one of the representative staining. (**c-d**). Immunofluorescence staining of TH and Lyve1 from contra-lateral and ipsilateral TDLNs from B16F10 subcutaneous tumor-bearing C57BL/6J mice. Photos shown one of the representative staining.

Further analysis revealed a significant accumulation of TH⁺ cells within the lymphatic structures of ipsilateral dcLNs in GSC005 tumor-bearing mice. A subset of these TH⁺ cells co-expressed CD3, suggesting the presence of CD3⁺TH⁺ T cells within the lymphatic compartment (**Fig. 2b**).

Solid tumors have been shown to invade and metastasize distant sites through both lymphatic dissemination^27^ and perineural spread^28^. This phenomenon has been well documented in multiple cancer types, including melanoma^29^ and breast cancer^30^. Consistent with these observations, the subcutaneous B16 melanoma model exhibited significant lymphatic expansion in ipsilateral TDLNs compared to contralateral TDLNs (**Fig. 2c**). Moreover, TH⁺ cells were prominently localized within these expanded lymphatic regions (**Fig. 2c**).

Collectively, these findings suggest that although TH⁺ signaling in the CNS and PNS differ in origin and composition, they share conserved functional roles in solid tumors, particularly in regulating lymphatic remodeling and facilitating tumor progression and dissemination.

### Therapeutic modulation of TH⁺ neurotransmitter receptors in tumors and TDLNs

TH⁺ neurons in the PNS are primarily components of the sympathetic nervous system (SNS), whereas in the CNS, TH⁺ neuronal populations include both dopaminergic and noradrenergic neurons^15^. Local and systemic cancer therapies may influence TH⁺ neuronal signaling by modulating the expression of neurotransmitter receptors in tumor cells as well as in non-tumor cells within TME and TDLNs.

To characterize neurotransmitter receptor expression in GBM, we first analyzed dopaminergic and adrenergic receptor profiles in patient tumor samples (**Fig. 3a-b**). The results revealed that both tumor and non-tumor cell populations exhibit relatively low expression of dopaminergic receptors, including the inhibitory receptor DRD4 and excitatory receptors DRD1 and DRD2 (**Fig.3a**). In contrast, noradrenergic signaling in the brain is primarily mediated through adrenergic receptors, including α- and β-adrenergic receptors. Among these, ADRB2 was identified as the predominant receptor expressed in tumor-infiltrating myeloid cells (**Fig.3b**).

**Fig. 3.**
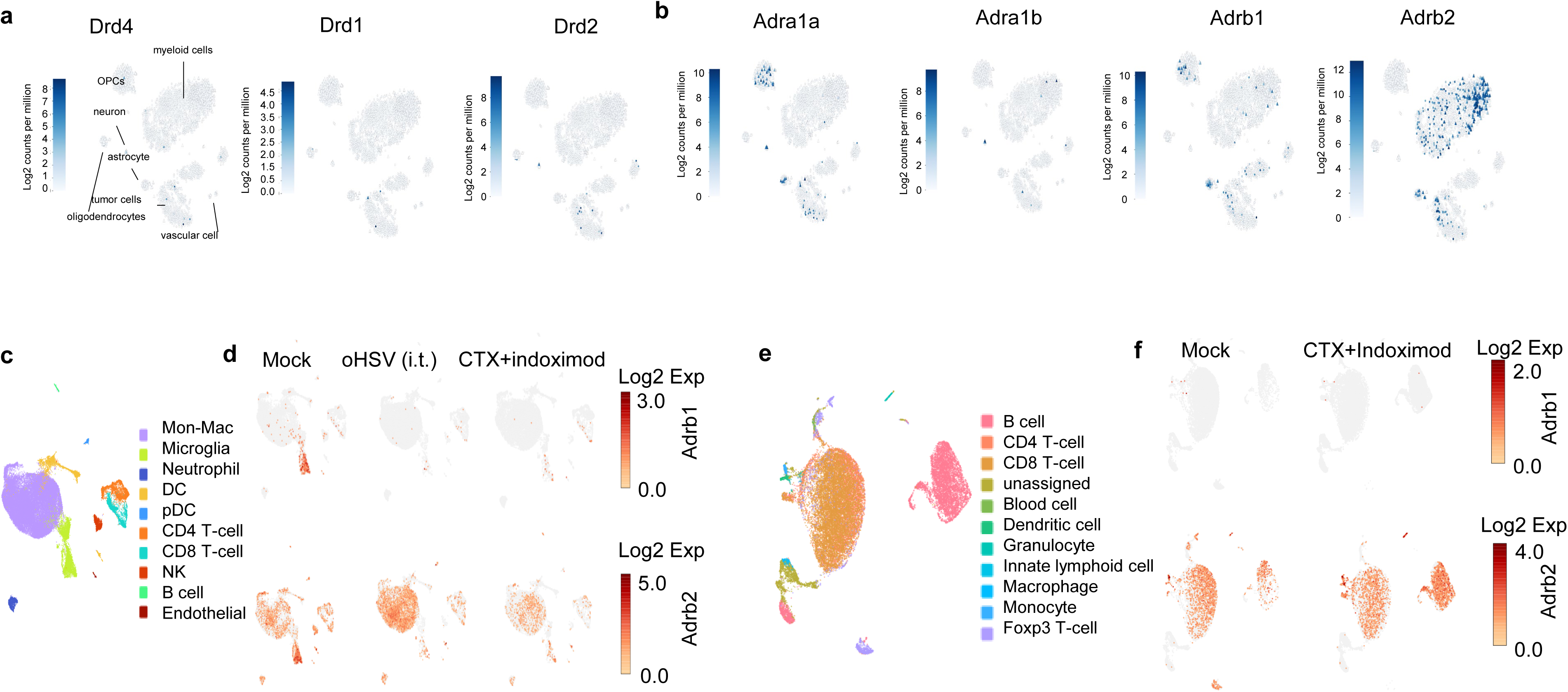
TH neurotransmitter receptors in human and murine GBM tumors. (**a-b**). Re-analysis of scRNA-seq data of TH neurotransmitter receptors in human GBM tumors. (**a**). Dopaminetic neurotransmitter receptors Drd4, Drd1 and Drd2 in human GBM. (**b**). Adrenenic receptors Adra1a, Adra1b, Adrb1 and Adrb2 in human GBM. (**c-d**). scRNA-seq analysis of adrenenic receptors Adrb1 and Adrb2 in GSC005 tumors treated with intra-tumor injection of oHSV or systemic treatment with cyclophosphamide(CTX) and indoximod. (**f-g**). scRNA-seq analysis of adrenenic receptors Adrb1 and Adrb2 in TDLNs of GSC005 tumors systemically treated with CTX and indoximod.

We next investigated how local and systemic therapies modulate TH signaling in tumor-infiltrating immune cells. Intratumoral administration of oHSV, which is known to rapidly enhance immune cell infiltration^21^, significantly altered adrenergic receptor expression (**Fig.3c-d**). Single-cell RNA sequencing (scRNA-seq) analysis demonstrated that oHSV treatment markedly upregulated β2-adrenergic receptor expression in myeloid cells—particularly in infiltrating macrophages—while concurrently downregulating β1-adrenergic receptor expression in microglia (**Fig. 3c-d**).

In contrast, systemic chemo-immunotherapy using cyclophosphamide (CTX) in combination with indoximod results in a distinct regulatory pattern. This treatment led to a downregulation of β2-adrenergic receptor expression in both myeloid cells and microglia within the tumor (**Fig. 3d**).

Since TDLNs for CNS tumors are located in peripheral tissues rather than within the brain, TH⁺ neuronal innervation in these sites differs from that within the tumor itself. Specifically, TH⁺ innervation of dcLNs, the TDLNs of CNS tumors, is mediated by the SNS. Analysis of adrenergic receptor expressions in dcLNs revealed the presence of β-adrenergic receptors across both lymphoid and myeloid cell populations (**Fig.3e-f**). Notably, systemic chemo-immunotherapy induced upregulation of β2-adrenergic receptor expression not only in T cells and myeloid cells but also in B cells (**Fig. 3f**).

### β-blockade enhances the efficacy of virotherapy

Given that the majority of tumor-infiltrating immune cells in brain tumors preferentially express β-adrenergic receptors rather than α-adrenergic receptors, we hypothesized that pharmacologic β-blockade could modulate TH⁺ neuronal signaling to enhance antitumor immunity. Consistent with this rationale, our previous study demonstrated that β-blockers can reprogram TH⁺ signaling in tumor-infiltrating myeloid cells and improve anti-tumor immune responses during intratumoral oHSV therapy in breast cancer^31^. Here, we investigated whether systemic administration of a β-blocker could enhance antitumor immune responses in GBM by targeting β-adrenergic signaling associated with TH⁺ neuronal innervation in both the tumor and TDLNs. In the GSC005 orthotopic GBM model, intratumoral injection of oHSV in combination with systemic β-blocker treatment (**Fig.4a**) significantly improved the survival of tumor-bearing mice (**Fig.4b**). This therapeutic benefit was accompanied by reduced tumor cell proliferation (**Fig. 4c**) and increased tumor cell death (**Fig.4d**). Immunohistochemistry analysis further revealed enhanced infiltration of CD8⁺ T cells in the combination treatment group (**Fig. 4e**), indicating potentiation of antitumor immune responses.

**Fig. 4.**
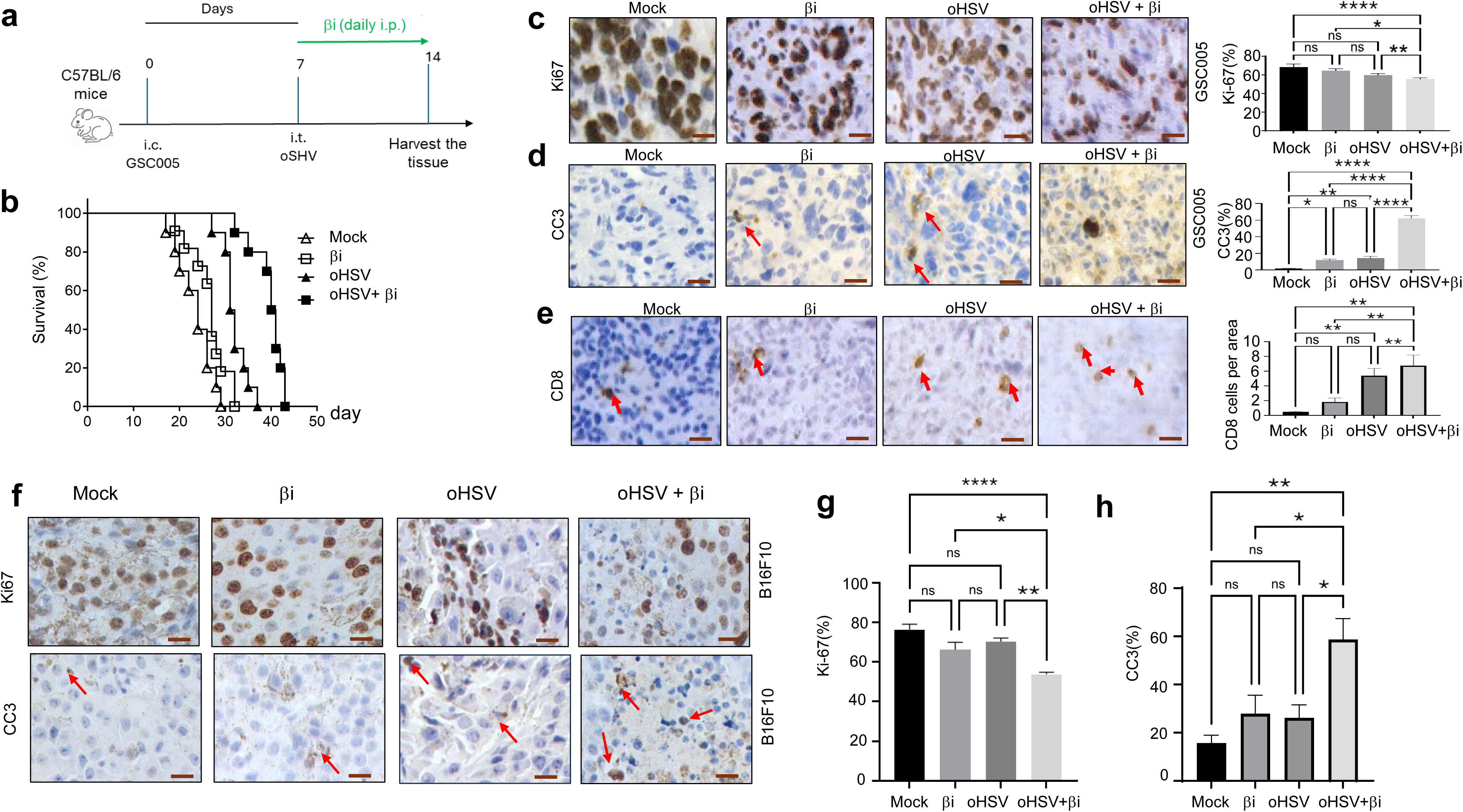
β-blocker improves anti-tumor effect of oHSV. (**a**). Diagram of the treatment of GSC005 tumor with oHSV and β-blocker (βi). (**b**).The Kaplan-Meier survival of GSC005 tumor-bearing mice treated with oHSV and βi. (n=10). *p*=0.026. (**c-e**). Immunohistochemistry staining of Ki-67 (**c**), cleaved caspase-3 (CC3, **d**) and CD8-T cells (**e**) of GSC005 tumor treated with oHSV and β-blocker. (n=5). \**p* < 0.05, \*\**p* < 0.01, \*\*\*\**p* < 0.0001, ns=no significance. (**f-h**). Immunohistochemistry staining of Ki-67, and cleaved caspase-3 (CC3) of subcutaneous B16F10 tumor treated with oHSV and β-blocker. (n=5). \**p* < 0.05, \*\**p* < 0.01, \*\*\*\**p* < 0.0001, ns=no significance.

To determine whether this effect extends to non-CNS tumors, we evaluated the combination therapy in a subcutaneous B16 melanoma model, which also exhibits TH⁺ neuronal innervation. Consistent with our findings in GBM, the combination of oHSV and β-blocker significantly suppressed tumor cell proliferation (**Fig. 4f-g**), increased tumor cell death (**Fig. 4f, 4h**), and CD8⁺ T cell infiltration (**sFig. 1**).

Collectively, these results demonstrate that β-adrenergic blockades enhance the efficacy of oncolytic virotherapy across both CNS and peripheral tumor models, likely through modulation of TH-mediated neuroimmune signaling.

### Inhibition of TH signaling enhances anti-tumor efficacy of intra-tumor injection of oHSV

In addition to functionally blocking SNS innervation through β-adrenergic antagonism, direct inhibition of TH activity represents an alternative strategy to suppress TH⁺ signaling within the tumor microenvironment. Pharmacologic inhibition of TH reduces catecholamine synthesis in TH⁺ cells, thereby attenuating downstream neuroimmune signaling. Several TH inhibitors, such as α-methyltyrosine (AMPT), have been evaluated in preclinical tumor models and shown to suppress tumor growth^32,33^, in part through modulation of cellular metabolism and induction of autophagy. To determine whether TH inhibition enhances the efficacy of oncolytic virotherapy, we evaluated the combination of a TH inhibitor with intratumoral administration of oHSV (**Fig.5a**). Consistently, the combination of oHSV with TH inhibition significantly suppressed tumor growth and increased tumor cell death compared to monotherapy (**Fig. 5b-e**). Furthermore, this combinatorial approach enhanced the infiltration of effector CD8⁺ T cells within the tumor microenvironment (**Fig.5f-g**), indicating improved antitumor immune activation.

**Fig. 5.**
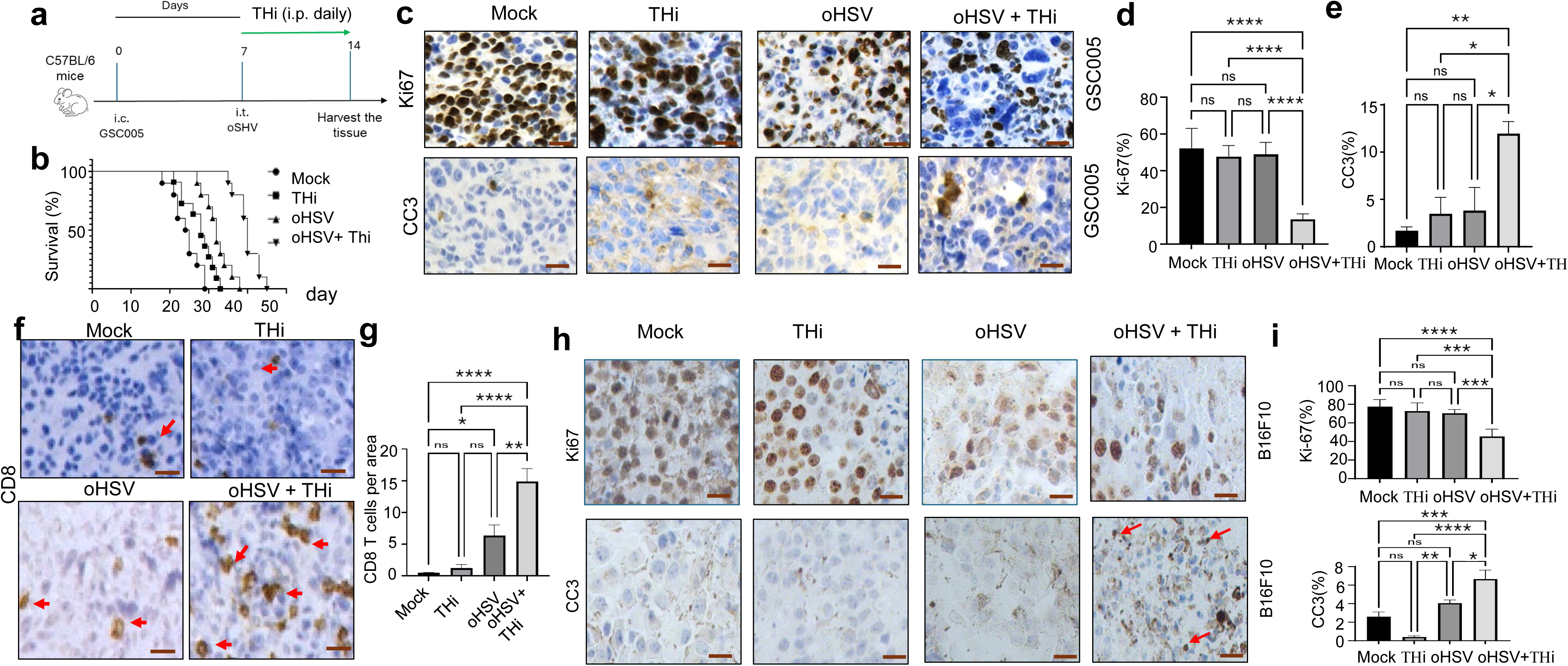
TH inhibitor improves anti-tumor effect of oHSV. (**a**). Diagram of the treatment of GSC005 tumor with oHSV and TH inhibitor (THi). (**b**). The Kaplan-Meier survival of GSC005 tumor-bearing mice treated with oHSV and THi. (n=10). *p*=0.018. (**c-g**). Immunohistochemistry staining of Ki-67(**c**, **d**), and cleaved caspase-3 (CC3, **c-e**), and CD8-T cells (**f-g**) of GSC005 tumor treated with oHSV and THi. (n=5). \**p* < 0.05, \*\**p* < 0.01, \*\*\*\**p* < 0.0001, ns=no significance. (**h-i**). Immunohistochemistry staining of Ki-67, and cleaved caspase-3 (CC3) of subcutaneous B16F10 tumor treated with oHSV and THi. (n=5). \**p* < 0.05, \*\**p* < 0.01, \*\*\*\**p* < 0.0001, ns=no significance.

To assess the generalizability of this strategy, the combination therapy was further evaluated in a subcutaneous B16 melanoma model. Similar antitumor effects were observed, including reduced tumor growth, increased tumor cell death (**Fig. 5h-i**), and enhanced CD8 T cells infiltration (**sFig.2**) Collectively, these findings demonstrate that direct inhibition of TH-mediated neurotransmitter synthesis represents an effective strategy to potentiate both local and systemic antitumor immune responses, particularly when combined with oncolytic virotherapy.

### Inhibition TH signaling modulates anti-tumor immune response in syngeneic 005 glioma model

To uncover the underlying mechanisms of improving anti-tumor immune response of combination TH signaling inhibition with intra-tumor oHSV therapy in syngeneic 005 glioma model, scRNA-seq assay was performed in the CD45 cells. Cells were clustered (**sFig.3** and **sFig.4**). Results indicated that the addition of TH inhibitor (α-Methyl-p-tyrosine significantly increased cytotoxic immune cells (CD8, NK and γδ T) (**Fig. 6a-b**) and inflammatory M1 macrophages (**Fig.6a-b**), while dramatically reduces immunosuppressive M2 macrophages (**Fig. 6a-b**). For tumor-infiltrated CD4 T cells, addition of TH inhibitor increases type-I IFN response (Irf7) and downregulate Apoe and Arg1 (**Fig.6c**). For cytotoxic CD8 T cells, addition of TH inhibitor increased T cell activation (CD69), cytotoxic molecules (Gzmb and Prf1), and downregulating Pdcd1 and Ctla4 (**Fig.6d**). Addition of TH inhibitor increases Fos and Ccl5 in activated NK cells (**Fig.6e**) and Fos and Ltb in γδ T cells (**Fig. 6f**) during oHSV therapy of 005 glioma. Since intra-tumor injection of oHSV significantly increases myeloid cells, especially macrophages^21^, addition of TH inhibitor significantly increases M1 macrophages (**Fig6g**) and decreases M2 macrophages (**Fig.6h**). Interestingly, addition of TH inhibitor induces feedback Adrb2 expression in M1 macrophages (**Fig.6g**), and downregulating Arg1 not only in M1 and M2 macrophages (**Fig.6h**) but also in other immune cells including CD4 (**Fig.6c**), NK(**Fig.6e**) and γδ T cells (**Fig. 6f**). The results indicate that TH inhibition could boost cytotoxic immune cells function but suppress immunosuppressive cells activity and induce feedback adrenergic signaling in tumor infiltrated immune cells.

**Fig. 6.**
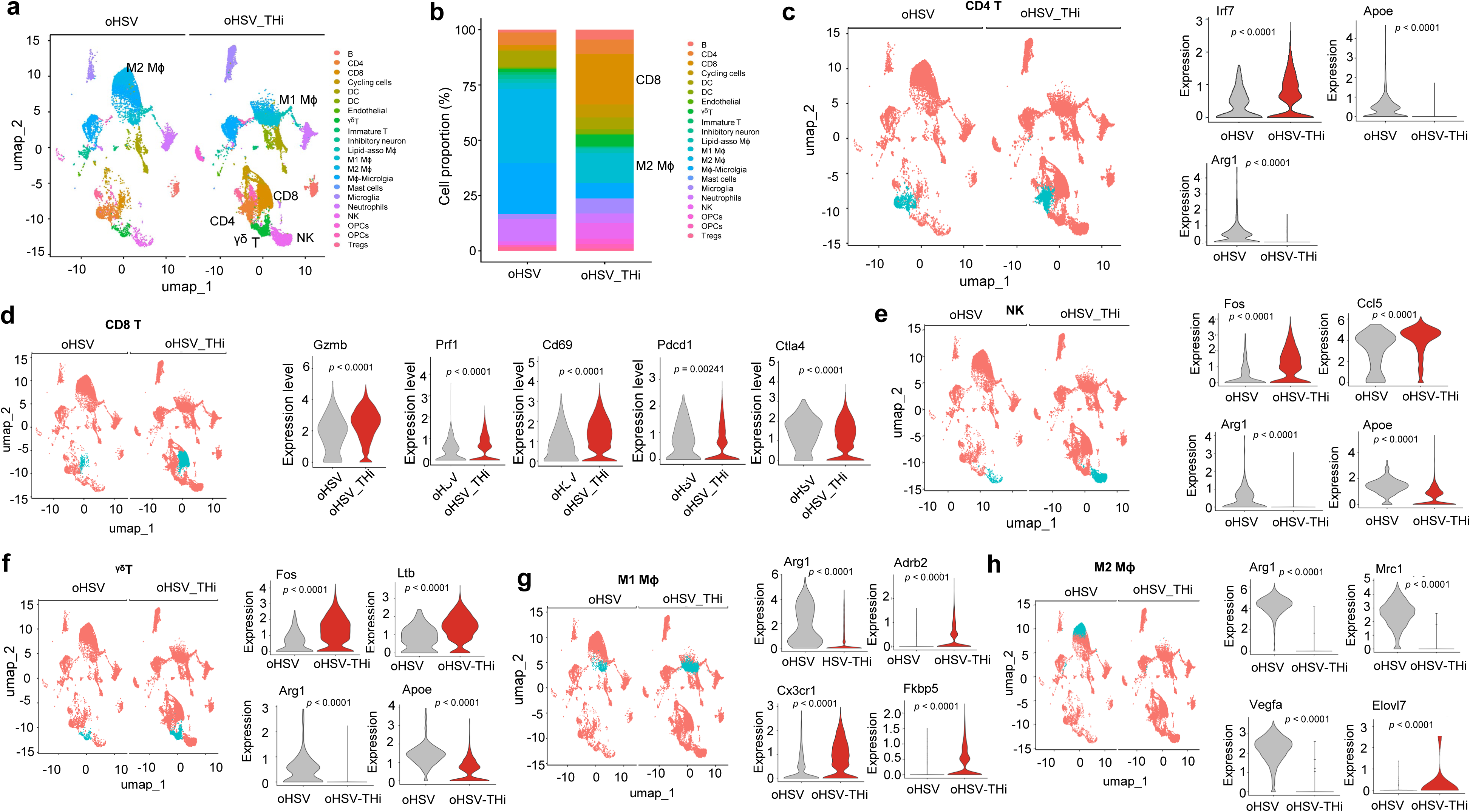
TH inhibitor modulates immune response in GBM during oHSV therapy. CD45+ cells from 005 tumor treated with oHSV or oHSV with THi were subjected to scRNA-seq analysis. **a**. UMAP of cell clusters in the 005 tumors. **b**. Immune cell clusters in the 005 tumors.**c**. UMAP of CD4 cells and vilion plot of Irf7, Apoe and Arg1 in CD4 cells. **d**. UMAP of CD8 T-cells and vilion plot of Gzmb, Prf1, Cd69, Pdcd1 and Ctla4 in CD8 T-cells. **e**. UMAP of NK cells and vilion plot of Fos, Ccl5, Arg1, and Apoe in NK cells. **f**. UMAP of γδ T-cells and vilion plot of Fos, Ltb, Arg1 and Apoe in γδ T-cells. **g**. UMAP of M1 macrophages (M1 Mɸ) and vilion plot of Arg1, Adrb2, Cx3cr1 and Fkbp5 in M1 Mɸ cells. **h**. UMAP of M2 macrophages (M2 Mɸ) and vilion plot of Arg1, Mrc1, Vegfra and Elovl7 in M2 Mɸ cells. The blue-green cluster in panel **b**-**h** represents the specific cell type analyzed.

## Discussion

In this study, we identify TH–mediated neuroimmune signaling as a previously underappreciated regulator of GBM progression and therapeutic response. By integrating single-cell RNA-sequencing, and functional therapeutic studies, our findings establish that TH⁺ neuronal and non-neuronal cells are not only present within the GBM TME, but also actively participate in shaping antitumor immunity in both the tumor and TDLNs. Importantly, we demonstrate that modulation of TH-associated signaling—either through β-adrenergic blockade or direct inhibition of TH enzymatic activity—significantly enhances the efficacy of oncolytic virotherapy, providing a strong rationale for targeting neuroimmune crosstalk as a therapeutic strategy in CNS tumors.

One of the key findings of this work is the identification of TH⁺ cells across multiple compartments of the GBM ecosystem, including neurons, astrocytes, and immune cells. While prior studies have largely focused on peripheral sympathetic innervation in solid tumors^11,31^, our data reveals that TH expression in the CNS tumor context is more heterogeneous, involving both classical neuronal populations and non-neuronal stromal and immune components. The presence of TH⁺ astrocytes and CD3⁺TH⁺ T cells^16^ suggests that catecholamine synthesis and signaling may occur locally within the tumor and lymphoid tissues, independent of long-range neuronal projections. This expands the current paradigm of neuroimmune regulation by indicating that neurotransmitter production in tumors is not solely neuron-derived, but may also arise from immune and glial compartments, thereby creating a complex and spatially distributed signaling network. Whether the CD3+TH+ cells includes the regulatory T cell population^16^ in the TDLNs or tumor need further investigated.

Our findings further highlight the importance of TH-mediated signaling in regulating lymphatic remodeling and immune activation in TDLNs^25^. The enrichment of TH⁺ cells in proximity to lymphatic vessels, along with the observed lymphangiogenesis in ipsilateral dcLNs^4,34,35^, suggests that neuroimmune interactions contribute to the structural and functional reprogramming of lymphatic niches during tumor progression. Given that dcLNs serve as primary sites for antigen presentation and T cell priming in GBM^34,36,37^, TH⁺ innervation may influence the quality and magnitude of systemic antitumor immune responses. The identification of CD3⁺TH⁺ T cells within lymphatic structures raises the intriguing possibility that T cells themselves may engage in autocrine or paracrine catecholamine signaling, potentially modulating their activation, differentiation, or exhaustion states. These observations align with emerging evidence in peripheral cancers, but extend the concept to CNS tumors, where lymphatic drainage and immune priming have only recently been appreciated.

Another important insight from this study is the dynamic regulation of neurotransmitter receptor expression in response to therapy. We show that oHSV treatment induces upregulation of ADRB2 in tumor-infiltrating myeloid cells, particularly macrophages, while systemic chemo-immunotherapy exerts distinct and sometimes opposing effects in the tumor and TDLNs. These findings suggest that therapeutic interventions can reshape the sensitivity of immune cells to neurogenic signals, thereby altering the balance between immunosuppressive and immunostimulatory pathways. The preferential expression of β-adrenergic receptors, especially ADRB2, in myeloid populations underscores the central role of catecholamine signaling in regulating myeloid cell function in GBM. Given that myeloid cells are key mediators of immunosuppression in the GBM TME^38,39^, targeting β-adrenergic signaling may represent an effective strategy to reprogram these cells toward a more pro-inflammatory phenotype.

Consistent with this hypothesis, our functional studies demonstrate that pharmacologic β-blockade significantly enhances the therapeutic efficacy of oHSV in both orthotopic GBM and subcutaneous melanoma models. The combination therapy not only improves survival but also reduces tumor proliferation, increases tumor cell death, and promotes infiltration of cytotoxic CD8⁺ T cells. These results suggest that β-adrenergic signaling acts as a negative regulator of antitumor immunity, and that its inhibition can synergize with virotherapy to amplify immune-mediated tumor clearance. Importantly, the consistency of these findings across CNS and non-CNS tumor models indicates that TH-mediated neuroimmune regulation may represent a conserved mechanism in solid tumors, despite differences in the origin and composition of TH⁺ innervation.

In addition to receptor-level modulation, we further demonstrate that direct inhibition of TH enzymatic activity using pharmacologic inhibitors such as AMPT can similarly potentiate the effects of oHSV therapy. By reducing catecholamine synthesis at its source, TH inhibition likely attenuates multiple downstream signaling pathways, including both adrenergic and dopaminergic axes. The observed enhancement of CD8⁺ T cell infiltration and tumor cell killing suggests that suppressing neurotransmitter production can relieve immunosuppressive constraints within the TME. Notably, TH inhibition may have broader effects beyond immune modulation, including alterations in tumor cell metabolism and stress responses, which could further contribute to therapeutic efficacy. Together, these findings highlight two complementary strategies—receptor blockade and neurotransmitter synthesis inhibition—for targeting TH-mediated signaling in cancer.

Despite these advances, several limitations of the current study should be acknowledged. First, while our data establishes strong associations between TH signaling and immune regulation, the precise molecular mechanisms by which catecholamines influence specific immune cell subsets remain to be fully elucidated. For example, the downstream signaling pathways activated by ADRB2 in myeloid cells and T cells in the GBM context require further investigation. Second, the relative contributions of neuronal versus non-neuronal sources of TH in driving tumor progression and immune modulation are not yet clearly defined. Genetic or cell type–specific ablation models will be necessary to dissect these contributions. Third, although our preclinical models provide robust evidence of therapeutic benefit, the translational relevance of these findings will need to be validated in human clinical settings, particularly given the complexity of neuroimmune interactions in patients.

In conclusion, our study provides compelling evidence that TH-mediated neuroimmune crosstalk is a critical determinant of tumor progression and therapeutic response in GBM. By demonstrating that both pharmacologic β-blockade and direct TH inhibition can enhance the efficacy of oncolytic virotherapy, we establish a novel therapeutic framework that integrates neural and immune targeting strategies. These findings not only advance our understanding of the neurobiology of cancer but also open new avenues for the development of combination therapies aimed at overcoming immunotherapy resistance in GBM and potentially other solid tumors.

## Acknowledgement

This work was supported by grants from the National Institute of Health (NIH) (R21NS130429 to BH), (NIH) (R61NS128191 to BH and BK), Alex Lemonade Stand Foundation Reach Grant (ALEX23-27891 to BH), and Paceline Foundation (MCG8451Tto BH).

## Author contributions

BH. designed experiments. KK, DZ, RK, TSJ, and BH performed experiments and analyzed the data. HD performed bulk mRNA-seq and scRNA-seq analysis. BH wrote the manuscript. BK and DHM revised the manuscript. All authors edited and reviewed the manuscript.

