## Supplemental Data for "Tyrosine hydroxylase–mediated neuroimmune crosstalk regulates antitumor immunity in glioblastoma during oncolytic herpes virotherapy"

**Supplementary Figure Legend**

**sFig.1 Immunohistochemistry staining of CD8 T cells in B16 tumor treated with oHSV and β-blcoker.** Subcataneous B16 tumors were treated with oHSV, or β-blcoker the combination. CD8 T-cell intiltration were analyzed by immunohistochemistry staining. (n=5). *$p < 0.05$, ****$p < 0.0001$, ns=no significance.

**sFig.2 Immunohistochemistry staining of CD8 T cells in B16 tumor treated with oHSV and TH inhibitor.** Subcataneous B16 tumors were treated with oHSV, or TH inhibitor the combination. CD8 T-cell intiltration were analyzed by immunohistochemistry staining. (n=5). *$p < 0.05$, ***$p < 0.005$, ****$p < 0.0001$, ns=no significance.

**sFig.3 Heatmap of cell annotation of scRNA-seq of CD45 cells from 005 tumors treated with oHSV or oHSV with THi.** GSC005 tumors were treated with oHSV, or oHSV with THi, or the combination. CD45 cells were subjected to scRNA-seq. Cell annotations were based on the top gene signatures.

**sFig.4 Heatmap of top 10 genes upregulated in immune cells from GSC005 tumors treated with oHSV or oHSV with THi.** GSC005 tumors were treated with oHSV, or oHSV with THi. CD45 cells were subjected to scRNA-seq. Top 10 genes upregulated in CD4, CD8, NK, γδ T-cell, M1 macrophage, M2 macrophage, dendritic cells and macrophage-microglia from oHSV or oHSV with THi treatment were displayed in the heatmap.

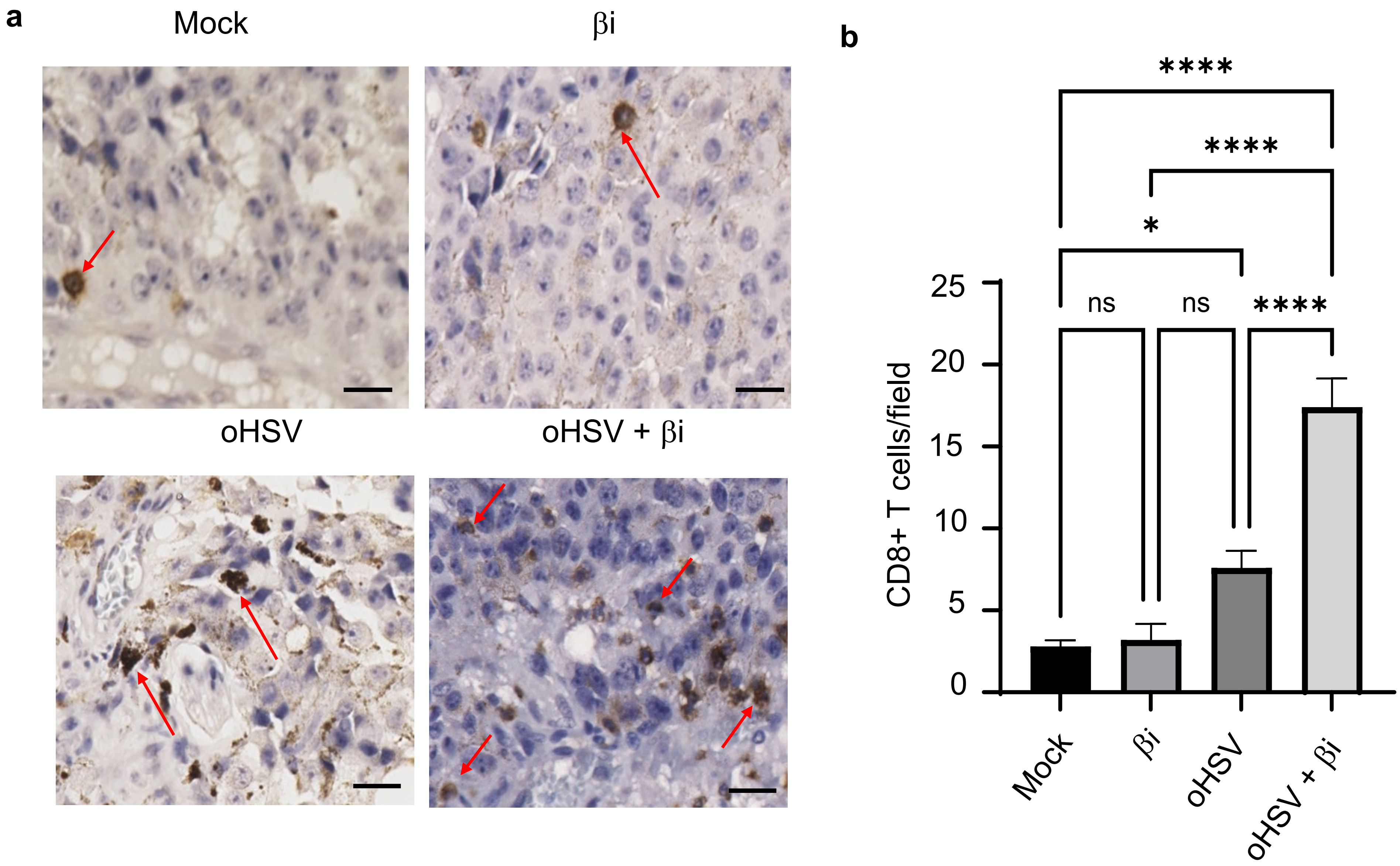
a
Mock
βi
oHSV
oHSV + βi
b
****
****
*
ns
ns
****
25
20
15
10
5
0
CD8+ T cells/field
Mock
βi
oHSV
oHSV + βi

sFig.1

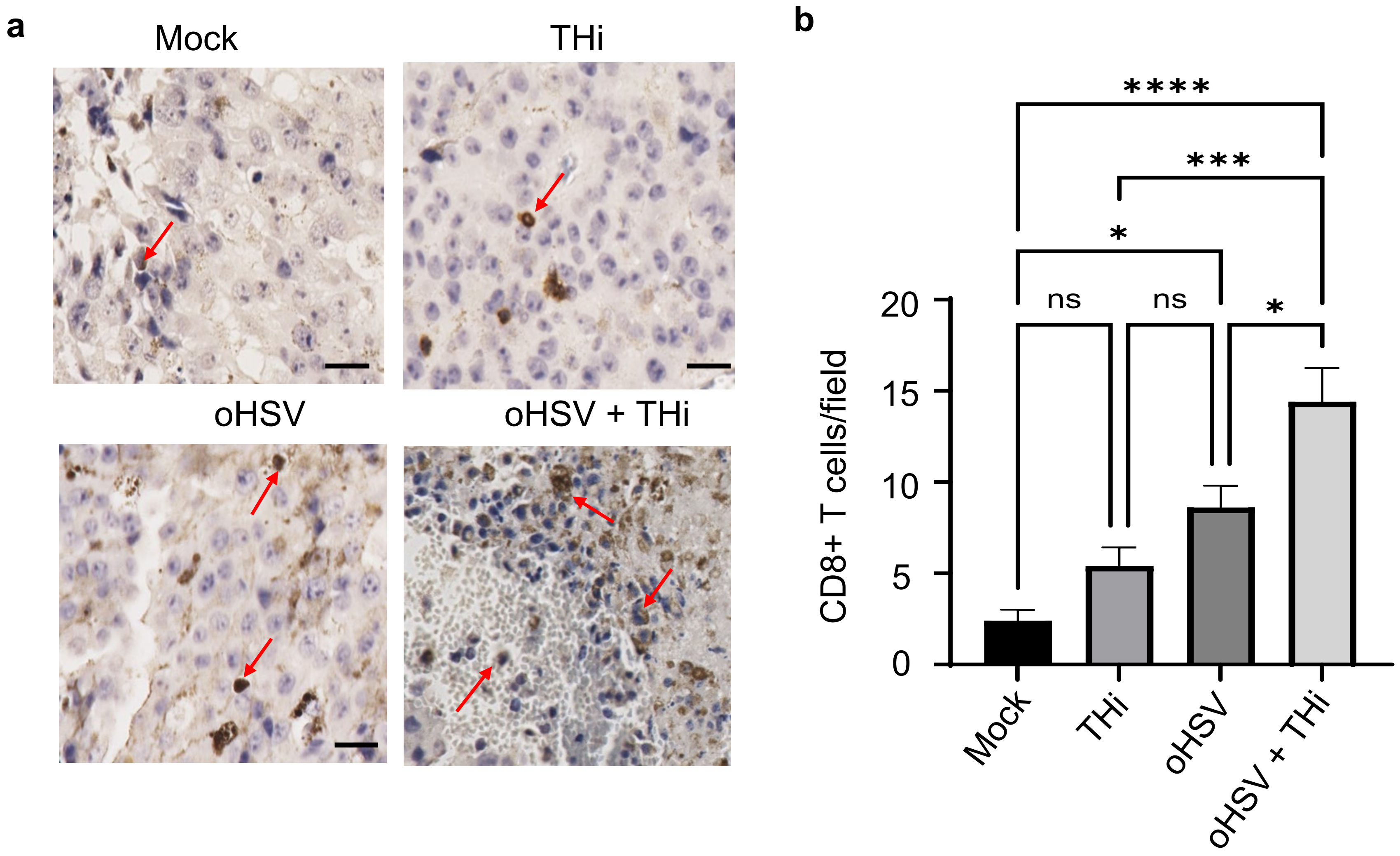
a
Mock
THi
oHSV
oHSV + THi
b
****
***
*
ns
ns
*
20
15
10
5
0
CD8+ T cells/field
Mock
THi
oHSV
oHSV + THi

**sFig.2**

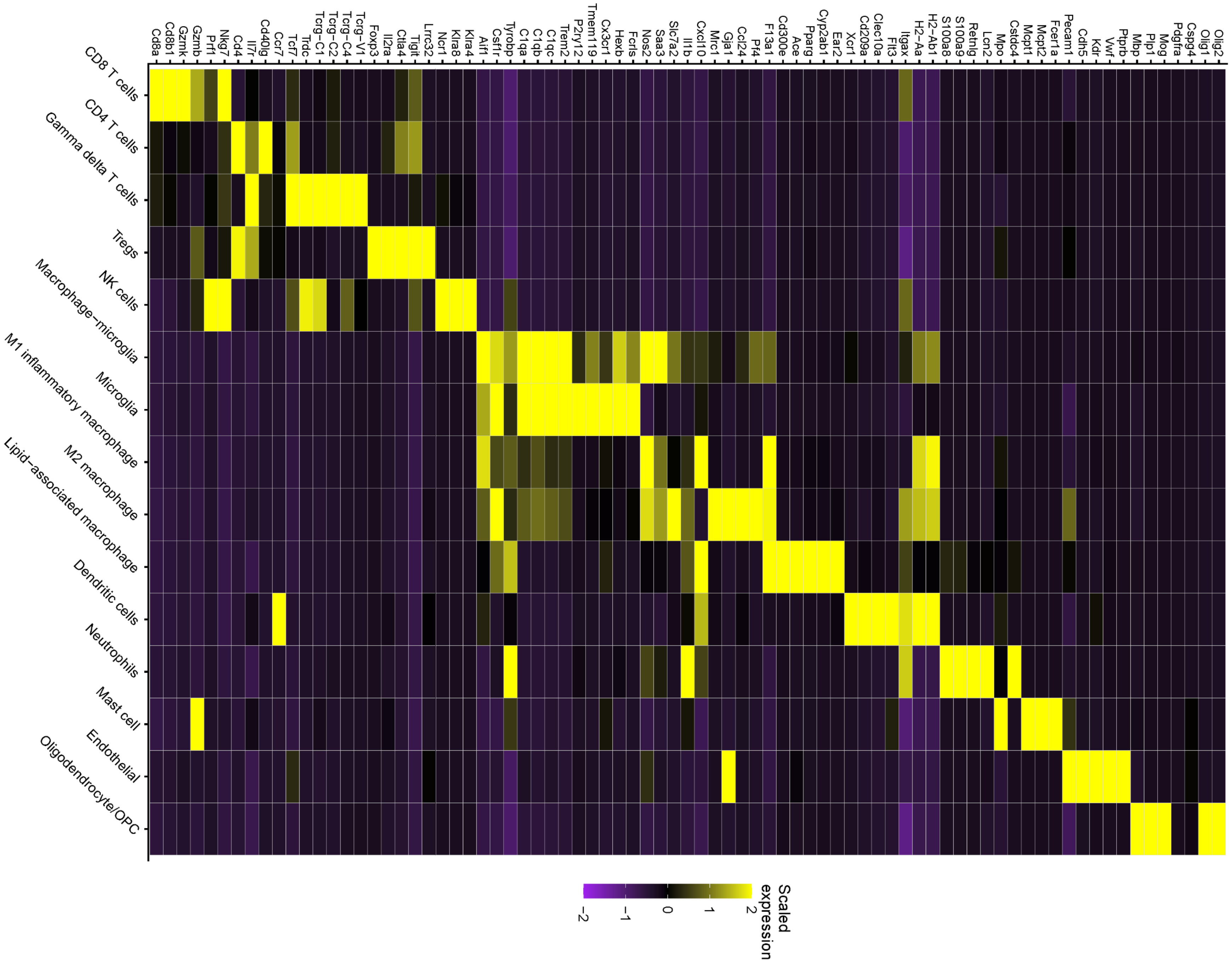
Cd8a
Cd8b1
Gzmk
Gzmb
Prf1
Nkg7
Cd4
Il7r
Cd40lg
Ccr7
Tcf7
Trdc
Tcrg−C1
Tcrg−C2
Tcrg−C4
Tcrg−V1
Foxp3
Il2ra
Ctla4
Tigit
Lrrc32
Ncr1
Klra8
Klra4
Aif1
Csf1r
Tyrobp
C1qa
C1qb
C1qc
Trem2
P2ry12
Tmem119
Cx3cr1
Hexb
Fcrls
Nos2
Saa3
Slc7a2
Il1b
Cxcl10
Mrc1
Gja1
Ccl24
Pf4
F13a1
Cd300e
Ace
Pparg
Cyp2ab1
Ear2
Xcr1
Cd209a
Clec10a
Flt3
Itgax
H2−Aa
H2−Ab1
S100a8
S100a9
Retnlg
Lcn2
Mpo
Cstdc4
Mcpt1
Mcpt2
Fcer1a
Pecam1
Cdh5
Kdr
Vwf
Ptprb
Mbp
Plp1
Mog
Pdgfra
Cspg4
Olig1
Olig2
CD8 T cells
CD4 T cells
Gamma delta T cells
Tregs
NK cells
Macrophage−microglia
Microglia
M1 inflammatory macrophage
M2 macrophage
Lipid−associated macrophage
Dendritic cells
Neutrophils
Mast cell
Endothelial
Oligodendrocyte/OPC
Scaled expression
−2
−1
0
1
2

**sFig3**

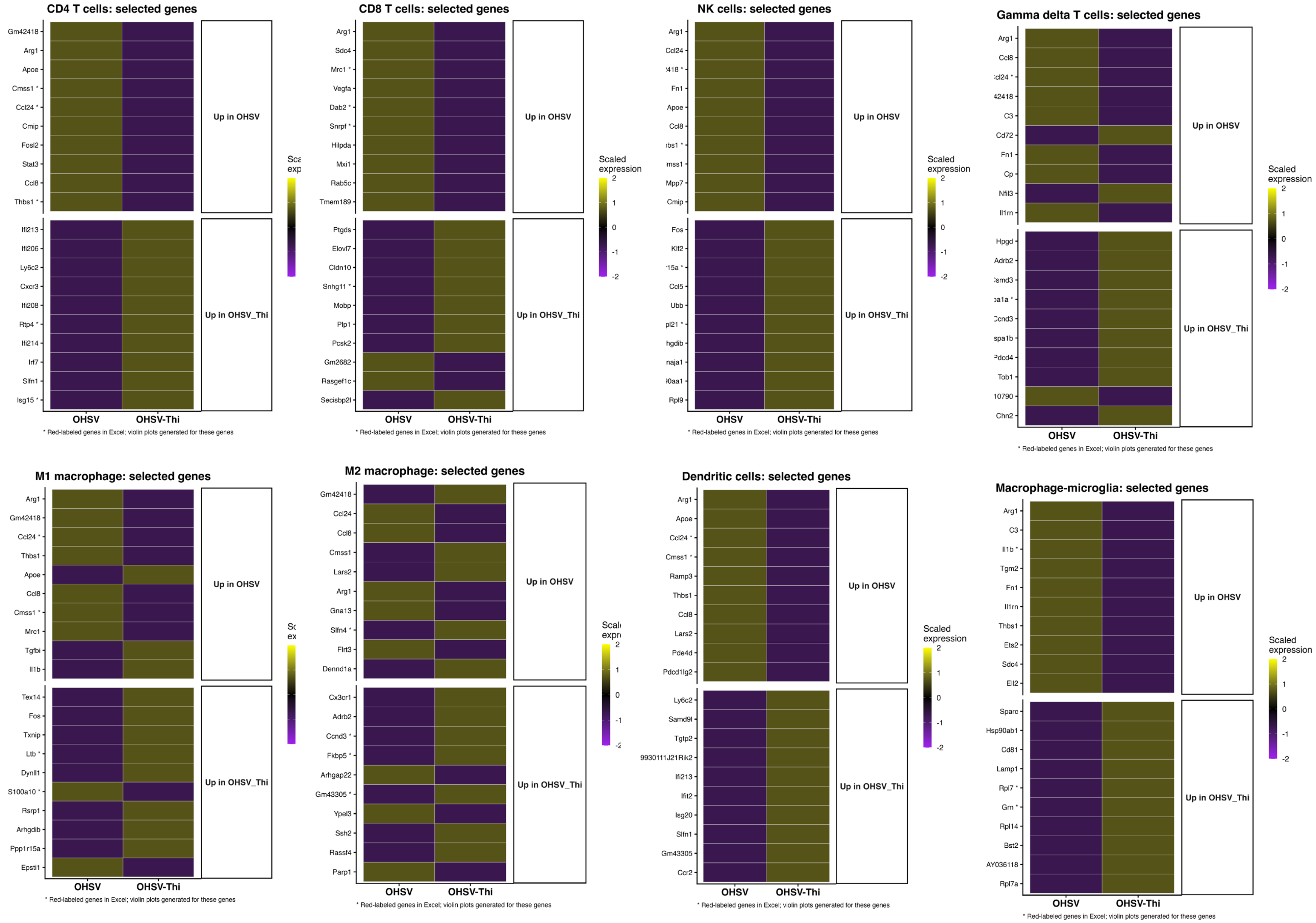
CD4 T cells: selected genes
Gm42418
Arg1
Apoe
Cmss1 *
Ccl24 *
Cmip
Fosl2
Stat3
Ccl8
Thbs1 *
Ifi213
Ifi206
Ly6c2
Cxcr3
Ifi208
Rtp4 *
Ifi214
Irf7
Slfn1
Isg15 *
Up in OHSV
Up in OHSV_Thi
OHSV
OHSV-Thi
* Red-labeled genes in Excel; violin plots generated for these genes
CD8 T cells: selected genes
Arg1
Sdc4
Mrc1 *
Vegfa
Dab2 *
Snrpf *
Hilpda
Mxi1
Rab5c
Tmem189
Ptgds
Elovl7
Cldn10
Snhg11 *
Mobp
Plp1
Pcsk2
Gm2682
Rasgef1c
Secisbp2l
Scaled expression
2
1
0
-1
-2
Up in OHSV
Up in OHSV_Thi
OHSV
OHSV-Thi
* Red-labeled genes in Excel; violin plots generated for these genes
NK cells: selected genes
Arg1
Ccl24
Fn1
Apoe
Ccl8
Mpp7
Cmip
Fos
Klf2
Ccl5
Ubb
Rpl9
Scaled expression
2
1
0
-1
-2
Up in OHSV
Up in OHSV_Thi
OHSV
OHSV-Thi
* Red-labeled genes in Excel; violin plots generated for these genes
Gamma delta T cells: selected genes
Arg1
Ccl8
C3
Cd72
Fn1
Cp
Nfil3
Il1rn
Hpgd
Adrb2
Tob1
Chn2
Scaled expression
2
1
0
-1
-2
Up in OHSV
Up in OHSV_Thi
OHSV
OHSV-Thi
* Red-labeled genes in Excel; violin plots generated for these genes
M1 macrophage: selected genes
Arg1
Gm42418
Ccl24 *
Thbs1
Apoe
Ccl8
Cmss1 *
Mrc1
Tgfbi
Il1b
Tex14
Fos
Txnip
Ltb *
Dynll1
S100a10 *
Rsrp1
Arhgdib
Ppp1r15a
Epsti1
Up in OHSV
Up in OHSV_Thi
OHSV
OHSV-Thi
* Red-labeled genes in Excel; violin plots generated for these genes
M2 macrophage: selected genes
Gm42418
Ccl24
Ccl8
Cmss1
Lars2
Arg1
Gna13
Slfn4 *
Flrt3
Dennd1a
Cx3cr1
Adrb2
Ccnd3 *
Fkbp5 *
Arhgap22
Gm43305 *
Ypel3
Ssh2
Rassf4
Parp1
Up in OHSV
Up in OHSV_Thi
OHSV
OHSV-Thi
* Red-labeled genes in Excel; violin plots generated for these genes
Dendritic cells: selected genes
Arg1
Apoe
Ccl24 *
Cmss1 *
Ramp3
Thbs1
Ccl8
Lars2
Pde4d
Pdcd1lg2
Ly6c2
Samd9l
Tgtp2
9930111J21Rik2
Ifi213
Ifit2
Isg20
Slfn1
Gm43305
Ccr2
Scaled expression
2
1
0
-1
-2
Up in OHSV
Up in OHSV_Thi
OHSV
OHSV-Thi
* Red-labeled genes in Excel; violin plots generated for these genes
Macrophage-microglia: selected genes
Arg1
C3
Il1b *
Tgm2
Fn1
Il1rn
Thbs1
Ets2
Sdc4
Ell2
Sparc
Hsp90ab1
Cd81
Lamp1
Rpl7 *
Grn *
Rpl14
Bst2
AY036118
Rpl7a
Scaled expression
2
1
0
-1
-2
Up in OHSV
Up in OHSV_Thi
OHSV
OHSV-Thi
* Red-labeled genes in Excel; violin plots generated for these genes

sFig4
